# Site-specific Halogenation of Peptides and Proteins using engineered Halogenase Enzymes

**DOI:** 10.1101/2022.07.19.500721

**Authors:** Barindra Sana, Ding Ke, Eunice Hui Yen Li, Timothy Ho, Jayasree Seayad, Hung A Duong, Farid J Ghadessy

## Abstract

We demonstrate novel in vitro halogenation of peptides by halogenase enzymes, and identify the (G/S)GW motif (HaloTryp Tag) as a preferred substrate. We further derive PyrH halogenase mutants showing improved halogenation of the HaloTryp Tag, both as a free peptide and when genetically fused to model proteins.

---

Numerous peptide-based therapeutics have recently been developed for diverse indications including diabetes, infectious disease and oncology^1-8^. Peptide drugs typically comprise non-canonical amino acids with chemical modifications that dramatically alter their physicochemical properties and efficacy^9-14^. Notably, halogenation can impart numerous desirable properties^15-23^including increased cell membrane and blood-brain-barrier^24-27^permeability, improved cytotoxicity^28^, enhanced target affinity^29^, selectivity gains, and reduced side effects. ^30, 31^

Halogenated peptides are typically made by incorporation of chemically synthesized amino acid analogs. The viability of post-synthesis *in vitro* peptide halogenation has not been widely explored, although post-translational peptide modification is common in living organisms. ^17, 32^ These modifications are carried out by specific enzymes that can potentially work *in vitro*. However, studies suggest that these enzymes have co-evolved orthogonally with specific peptide substrates, limiting their application as generic peptide halogenation tools. ^20, 33^ Tryptophan halogenases (TH) catalyse halogenation of the free amino acid tryptophan, both *in vitro* and in living organisms.^20, 34, 35^

TH family members are classified based on their regiospecificty towards tryptophan (5, 6 or 7 positons of the indole ring). These have been engineered extensively to alter regioselectivity and improve activity towards non-natural chemical substrates. Here, we show for the first time that TH enzymes can additionally halogenate C-terminal tryptophan residues of synthetic peptides in a context-dependent manner. We further improve this novel activity by structure guided engineering and utilise these TH variants to site-specifcally halogenate model proteins.

The TH enzymes PyrH (T5H), SttH and ThHal (T6H), PrnA and RebH (T7H) were first assayed for chlorination of a panel of short 2-4 mer peptides comprising only tryptophan (W) and glycine (G) residues. PyrH, SttH and ThHal chlorinated peptides with tryptophan at the C-terminus (Table 1). The T7H enzymes PrnA and RebH did not produce any detectable chlorinated products. Interestingly, only mono-chlorinated products were observed, even for the peptides containing two tryptophan residues. Chlorination of WW and GW dipeptides was observed but not WG, suggesting that only the C-terminal tryptophan residue was accessible to the substrate binding site.

**Table 1.**
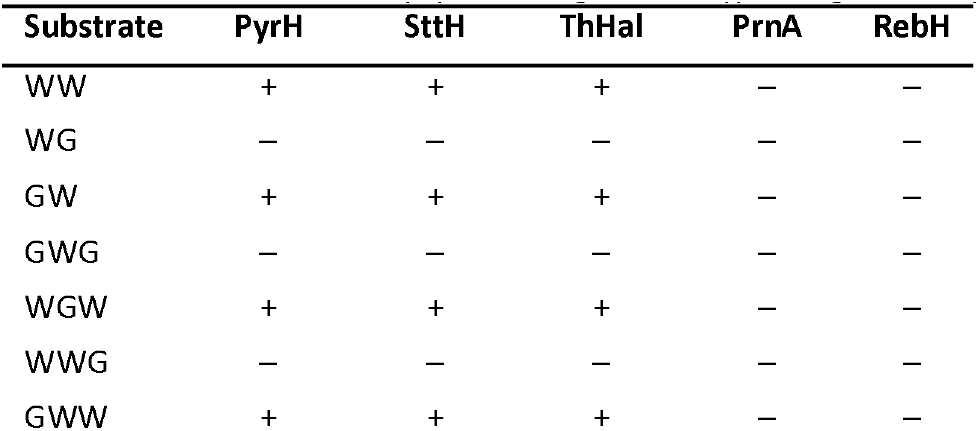
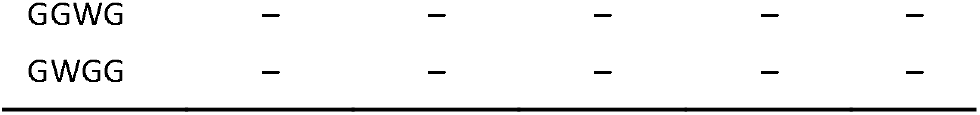
Chlorination of short peptides using five wild type halogenase enzymes.

Next, we expanded the panel of screened dipeptide substrates to include charged, nucleophilic and bulkier amino acid residues adjacent to the C-terminus tryptophan. PyrH exhibited very high activity on SW, GW and AW dipeptides, showing 83%, 55% and 50% chlorination respectively (Figure 1). SttH and ThHal also showed higher activity on these peptides. Peptides with bulkier or charged amino acid residue next to the C-terminal W showed reduced halogenation. All three enzymes were able to brominate the dipeptides, although with lower efficiencies. The most efficient enzyme PyrH was able to brominate 25% of the GW and 17% of the SW peptide.

**Figure 1.**
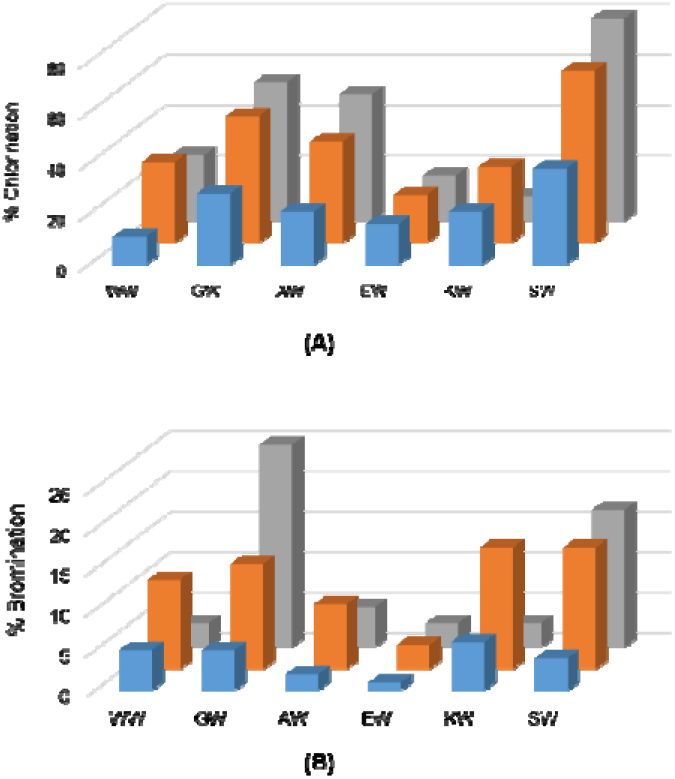
A) Screen for chlorination and B) bromination of indicated dipeptides using ThHal (blue), SttH (orange) and PyrH (grey) enzymes.

Tripeptides comprising the permissive SW, GW and AW motifs extended by either a bulky, nucleophilic, or charged amino acid at the N-teminus were next investigated. The halogenases were largely inefficient at chlorinating the tripeptides SSW, GSW, AAW or GAW, and almost inactive for their bromination (Figure 2). PyrH-catalyzed chlorination dropped from 83% for SW dipeptide to 16% and 10% for GSW and SSW tripeptides respectively. Activity on the AAW (9%) and GAW (1%) tripeptides was also markedly lower than for the AW dipeptide (50%). In contrast, chlorination and bromination of the tripeptide GGW (50%) was comparable to GW dipeptide (55%). Highest activity was seen for the SGW tripeptide, with 58% chlorination achieved by PyrH. Incorporation of charged amino acids reduced efficiency in all sequence contexts.

**Figure 2.**
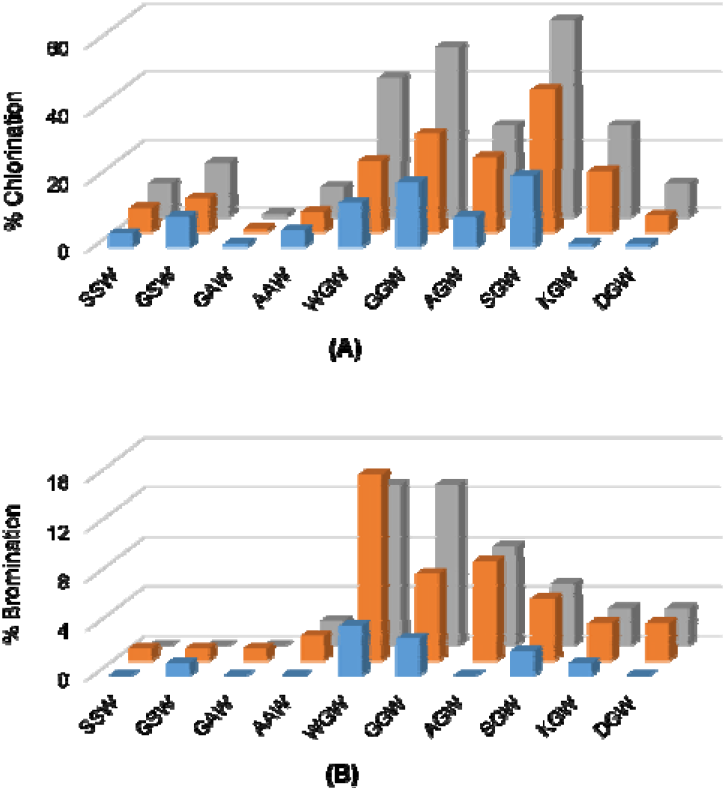
A) Screen for chlorination and B) bromination of indicated tripeptides using ThHal (blue), SttH (orange) and PyrH (grey) enzymes.

The effect of peptide length on PyrH-catalyzed chlorination was tested next by sequential N-terminal extension of the GGW and SGW tripeptides with glycine (Figure 3). All peptides were efficiently converted in comparison to the GW dipeptide and GGW /SGW tripeptides. The longer G_5_W and G_3_SGW peptides were next enzymatically chlorinated and purified at preparative scale. 1H NMR confirmed tryptophan C5 as the sole halogenation site (Figure S1 – S4), in agreement with the regiospecificity of PyrH.

**Figure 3.**
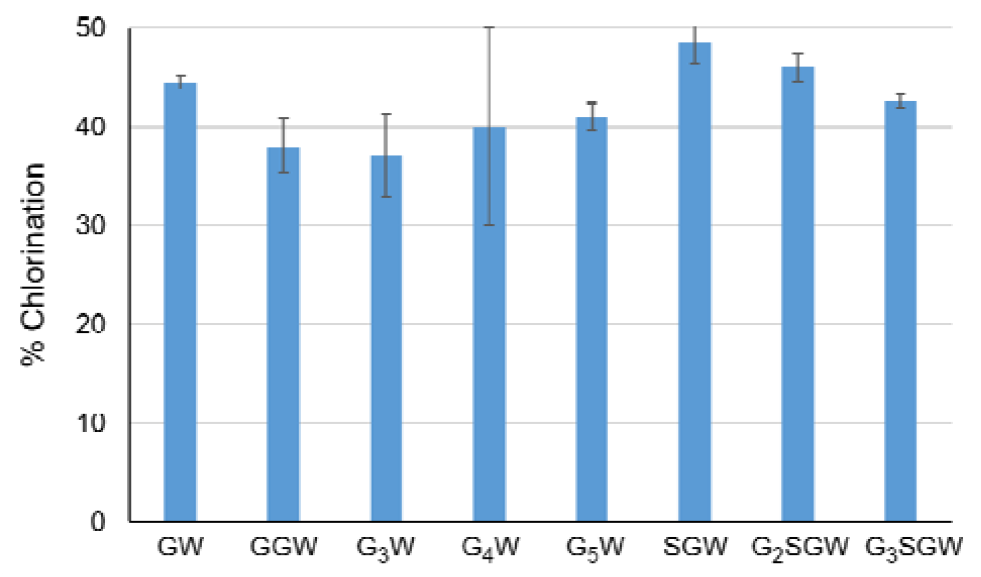
Chlorination of indicated peptides using PyrH. Values represent average ± SD (n=2)

The PyrH substrate binding pocket differs notably from PrnA and RebH, resulting in tryptophan adopting a different bound conformation^36^. Differences include both an α-helical region in the T7H enzymes (F_458_-N_464_ in RebH and F_447_-N_453_ in PrnA) and a short loop insertion (T_343_-F_438_ in RebH and N_444_-S_448_ in PyrH) that potentially limit optimal peptide binding by active site occlusion. No similar structural elements are present in PyrH, likely rendering its active site more accessible to the larger non-cognate peptide substrates (Figure 4). We hypothesized that further opening up of the PyrH substrate binding site could enhance halogenation efficiency. We therefore deleted four amino acids (A_145_-V_148_) within a proximal unstructured loop region (F_144_-Y_166_) to yield variant PyrH -dASQV. This enzyme showed between 10 to 30% increased activity over the wild-type PyrH on peptide substrates (Figure 5). We also introduced a conservative mutation, Q160N, to both increase active site accessibility and further accommodate residues preceding the C-terminal tryptophan (Figure 4). This variant (PyrH-Q160N) showed ≥50% higher activity over PyrH for chlorination of GGW and G_5_W peptides (Figure 5). The relative binding affinity (Km) of PyrH-Q160N was 34% higher than PyrH for the GGW peptide. The turn-over number (k_cat_) was similar for both variants, indicating that the modest increase in activity of PyrH-Q160N arose from improved substrate binding.

**Figure 4.**
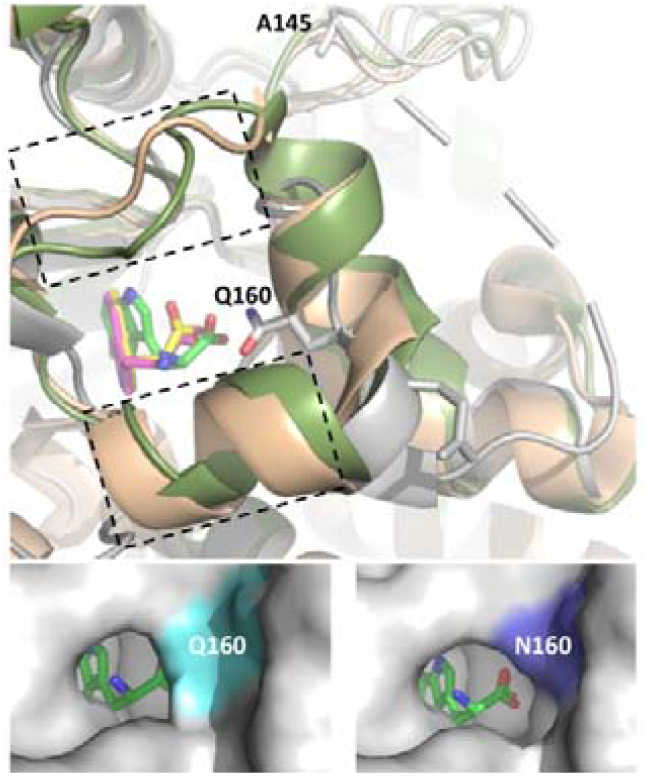
Top Panel: Structural overlay of PyrH (grey, bound tryptophan in green), PrnA (wheat, bound tryptophan in magenta) and RebH (green, bound tryptophan in yellow). Boxed region denotes a-helical region (F458-N464 in RebH and F447-N453 in PrnA) and loop region (T343-F438 in RebH and N444-S448 in PyrH) that is absent in PyrH. The PyrH residues Q160 (mutated to N in Q160N variant) and A145 (deleted in PyrH – d ASQV) are highlighted. Note that S146, Q147 and V148 also deleted in this mutant are not resolved in crystal structure. Bottom panels: Modelling of the PyrH Q160N mutation with bound tryptophan highlights cavitation of the active site region enabling access to larger peptidic substrates with C-terminal tryptophan residue. Images generated based on structures 2WEU^36^ (PyrH), 2OA1^37^(RebH) and 2ARD^38^ (PrnA).

**Figure 5.**
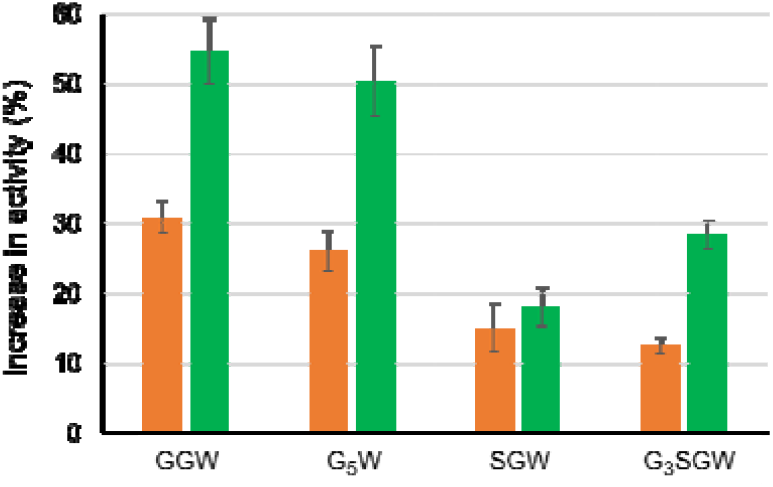
Peptide chlorination using PyrH-dASQV (orange) and PyrH-Q160N (green) for indicated peptides. Values indicate % increase in activity over PyrH ± SD (n=2).

We next recombinantly expressed and purified 3 model proteins, eGFP (30 KDa), Stoffel fragment of Taq DNA polymerase (64 KDa) and SpyCatcher (13 KDa) with genetically encoded C-terminal GGW or SGW tags (Table S1). C-terminal halogenation by PyrH and its two mutant variants was observed (Figure 6). This approached 90% conversion for the eGFP-GGW substrate using the PyrH mutants. As before, PyrH-Q160N was generally the most active (Figure 4). Lower halogenation efficiencies of the Stoffel and SpyCatcher proteins (∼40% using PyrH-Q160N) was observed, which may arise from steric hindrance between the halogenase and the substrate protein. Further iteration of linker lengths between these proteins and the (G/S)GW tag is warranted to address this possibility.

**Figure 6.**
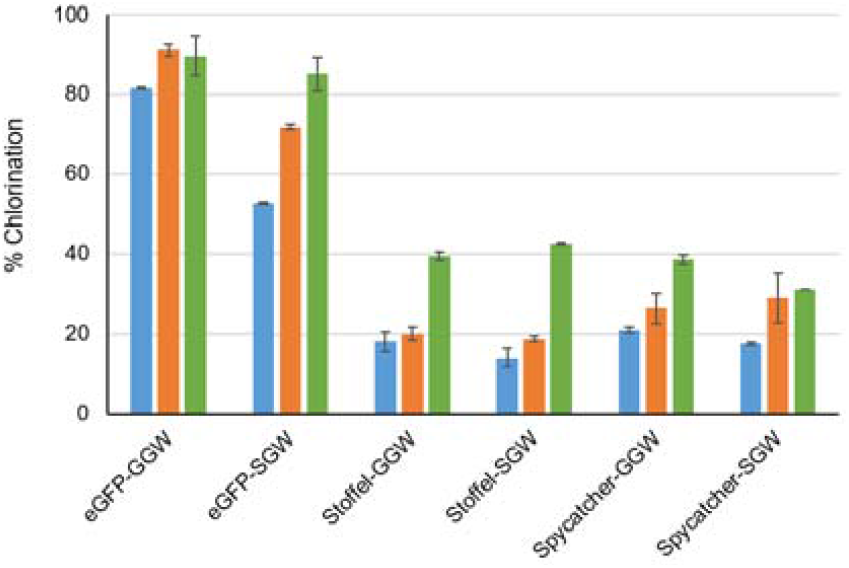
Chlorination of indicated proteins using WT PyrH (blue), PyrH-dASQV (orange) and PyrH-Q160N (green) mutants. Values represent average ± SD (n=2)

## Conclusions

We have demonstrated *in vitro* C-terminal enzymatic halogenation of a range of peptides. Screening yielded the optimal (G/S)GW motif, which we term the HaloTryp Tag. In combination with the rationally designed PyrH-Q160N mutant, the HaloTryp Tag facilitated efficient (40 to 90% conversion) site-specific halogenation of several model proteins. Post-translational labeling using the HaloTryp Tag could be used downstream of co-translational labeling methodologies^39^ to expand chemical diversity and potentially introduce novel physico-chemical properties.

