## Supplemental File 1 for "Site-specific Halogenation of Peptides and Proteins using engineered Halogenase Enzymes"

**MATERIALS AND METHODS**

**Cloning and enzyme engineering:** The halogenase genes were codon-optimized for recombinant expression in *E. coli*, and cloned in between NdeI and XhoI restriction sites of pET22b vector by infusion cloning. The construct was transformed into chemically competent DH5α cells, and the sequence of the construct was confirmed by DNA sequencing.

The PyrH Q160N mutant was made by PCR-based site directed mutagenesis using 5'- AGC ACC TTG GCA GAG AAT CGT GCA CAA TTC CCT TAC -3' and 5'- GTA AGG GAA TTG TGC ACG ATT CTC TGC CAA GGT GCT -3' primers. The PCR product was DpnI digested and transformed into chemically competent DH5α cells. The dASQV mutant was developed by inverse PCR of the wild type PyrH construct (using 5'- GAC GAA TCC CTG GGT CGT AGC ACC TTG -3' and 5'- GAA TAA TGA TCC ATC AAG CAT ACG TGG TGC GCG TT -3' primers) followed by DpnI digestion, gel purification and intramolecular ligation of the PCR product. The intramolecular ligation product was transformed into chemically competent DH5α cells. The correct mutations were confirmed by DNA sequencing.

**Enzyme expression and purification:** The correct plasmid constructs were transformed into *E. coli* BL21(DE3) competent cells following manufacturer’s protocol (NEB) and grown overnight at 37 °C in LB-agar plates containing suitable antibiotic. Overnight cultures were prepared by inoculating single colonies in antibiotic-supplemented LB broth and grown at 37 °C with constant shaking. For overexpression, the overnight cultures were diluted 100-fold in the same media, grown at 37 °C until the OD600 reached to 0.5, and induced with 0.2 mM IPTG. The cells were harvested after culturing overnight at 25 °C, and stored at –­­80 °C.

The cells were lysed by sonication and the proteins were purified using Ni-NTA column (GE healthcare) following standard protocols. The purified proteins were concentrated and buffer exchanged into 50 mM phosphate buffer pH 7.2, using 10 kDa MWCO Amicon Ultra centrifugal filters. Concentration of the purified proteins was determined by OD_280_ using Nanodrop and their purity was analyzed by SDS-PAGE.

**Halogenation reactions:** For peptide halogenation 2.5 mM substrate was treated with 10 µM enzyme and 50 mM NaCl/NaBr in the presence of the cofactors 10 µM FAD, 2 mM NADH and 30 µM flavin reductase enzyme RebF, 20 mM glucose and 5 units glucose dehydrogenase enzyme. The reaction was done in 10 mM phosphate buffer (pH = 7.2), overnight at room temperature with constant mixing. The enzymes were inactivated by heating at 95 °C for 10 minutes and removed by centrifugation at 13,500 rpm for 10 minutes. The supernatant was analysed by LC-MS, and the halogenated and non-halogenated products were quantified from the Area Under Curve.

For kinetic study, various concentrations (0.5 – 50 mM) of GGW peptide were halogenated by PyrH-WT and PyrH-Q160N enzymes following the above protocol. The reactions were stopped after 15 minutes by inactivating the enzymes at 95 °C for 10 minutes and the precipitates were removed by centrifugation at 13,500 rpm for 10 minutes. The supernatant was analysed by LC-MS. The amounts of halogenated products were quantified in terms of Area Under Curve, and used for calculating the relative reaction rate. The substrate concentrations and the relative reaction rates were used to make Lineweaver-Burk plot, which was used to calculate relative kinetics parameters.

For protein chlorination, 86 µM protein was treated with 10 µM enzyme and 50 mM NaCl in the presence of the cofactors 10 µM FAD, 2 mM NADH and 30 µM flavin reductase enzyme RebF, 20 mM glucose and 5 units glucose dehydrogenase enzyme. The reaction was done in 10 mM phosphate buffer (pH = 7.2), for 3 hours at room temperature with constant mixing. Precipitates were removed by centrifugation at 13,500 rpm for 10 minutes. To cleave the halogenated C-terminus, the supernatant was mixed with PreScission protease (10 unit/ml final concentration) and incubated overnight at 4 °C with constant mixing. The reaction mixtures were heated to 95 °C for 10 minutes, precipitates were removed by centrifugation at 13,500 rpm for 10 minutes. The supernatant was analysed by LC-MS, and the halogenated and non-halogenated cleaved C-terminus was quantified from the Area Under Curve.

**Detection of halogenation position:** The halogenation position was detected in the chlorinated products of two peptides G_5_W and G_3_SGW. The peptides were chlorinated in preparative scale by scaling up the above reaction and the chlorinated products were purified by semi-preparative HPLC. The halogenation site was identified by ^1^H NMR.

**Analytical methods:** Chemicals and anhydrous solvents were obtained from Sigma Aldrich and were used without further purification. Spectroscopic grade solvents were purchased from Sigma Aldrich. NMR spectra were recorded on Bruker Advance III 400MHz spectrometer in MeOD-d4. Data are reported in the following order: chemical shifts are given (δ); multiplicities are indicated as s (singlet), d (doublet), t (triplet), q (quartet) and m (multiplet). High-resolution mass spectra (HRMS) were recorded on an Agilent ESI-TOF mass spectrometer at 3500 V emitter voltage. Exact m/z values are reported in Daltons.

For analytical HPLC, 20 μL of crude mixture was injected onto an Agilent EclipsePlus C18 analytical column (1.8 μ packing, 2.1 mm × 50 mm). Gradient starting conditions of 10% (v/v) MeCN/H_2_O (plus 0.1 % (v/v) HCOOH) were held for 1 min before development to 95 % (v/v) MeCN/H_2_O over 3 min prior to re-equilibration to starting conditions over 2 min. Flow rates and column temperature were kept constant at 0.4 mL min^-1^ and 25 ^o^C respectively. UV absorbance was detected at 254 nm and 280 nm throughout.

For semi-preparative HPLC, 900 μL of solution containing crude mixture dissolved in H_2_O/MeCN was injected onto a Phenomenex Jupiter semi-preparative C12 HPLC column (4 μ packing, 250 x 10 mm). Starting conditions of 50 % (v/v) MeCN/H_2_O (plus 0.05 % (v/v) TFA) were held for 2 min. prior to development to 90 % (v/v) MeCN/H2O over 15 min. 95 % (v/v) MeCN/H_2_O then held for 3 min. prior to re-equilibration of starting conditions over 3 min. Flow rates were kept constant at 5 ml min^-1^. UV absorbance was detected at 280 nm throughout.

**SUPPLEMENTARY FIGURES**

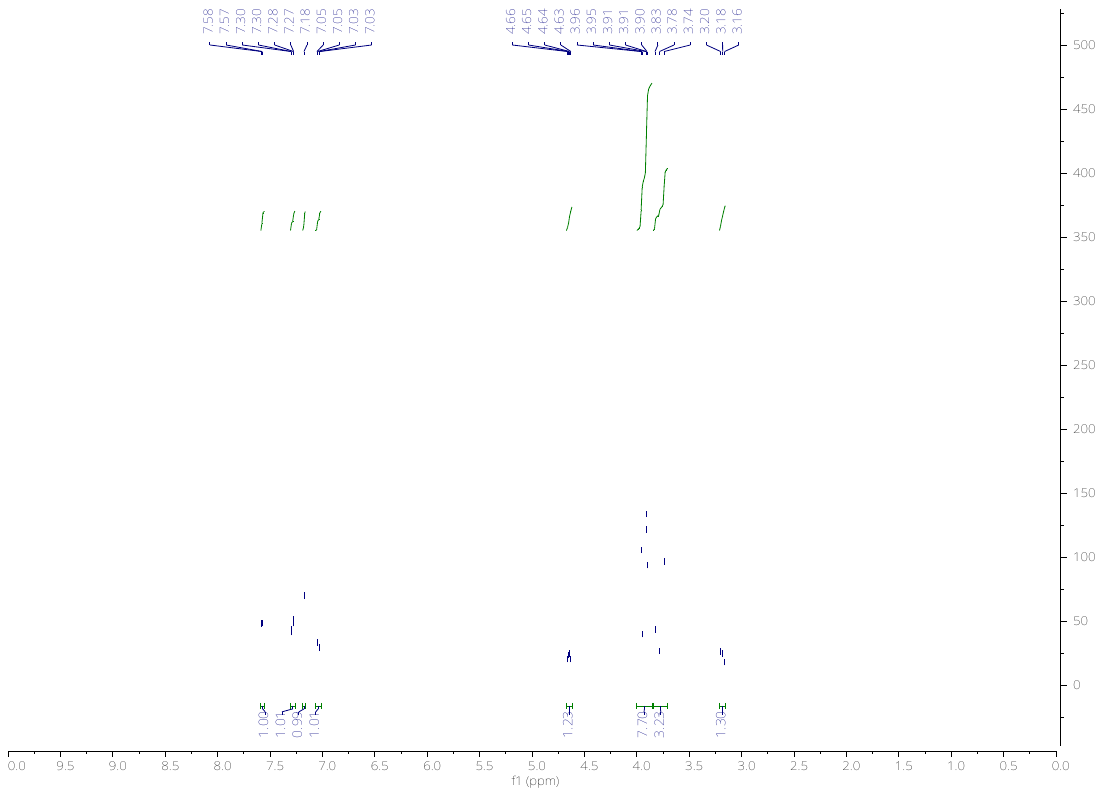

**Figure S1.** ^1^H Spectra of chlorinated G_5_W.

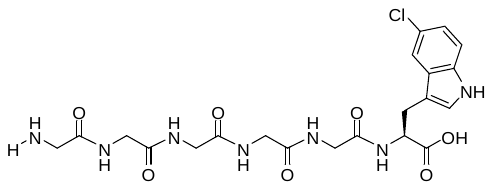

**Figure S2.** Chemical structure of chlorinated G_5_W.

1H NMR (400 MHz, CD_3_OD): δ 7.58 (d, J = 1.7 Hz, 1H), 7.29 (dd, J = 8.6, 0.5 Hz, 1H), 7.18 (s, 1H), 7.04 (dd, J = 8.6, 2.1 Hz, 1H), 4.64 (dd, J = 7.7, 5.1 Hz, 1H), 3.99 – 3.87 (m, 8H), 3.84 – 3.70 (m, 3H), 3.21 – 3.16 (m, 1H).

HRMS (ESI+): m/z calcd for C_21_H_26_ClN_7_O_7_ [M+H]^+^ 524.1655, found 524.1655.

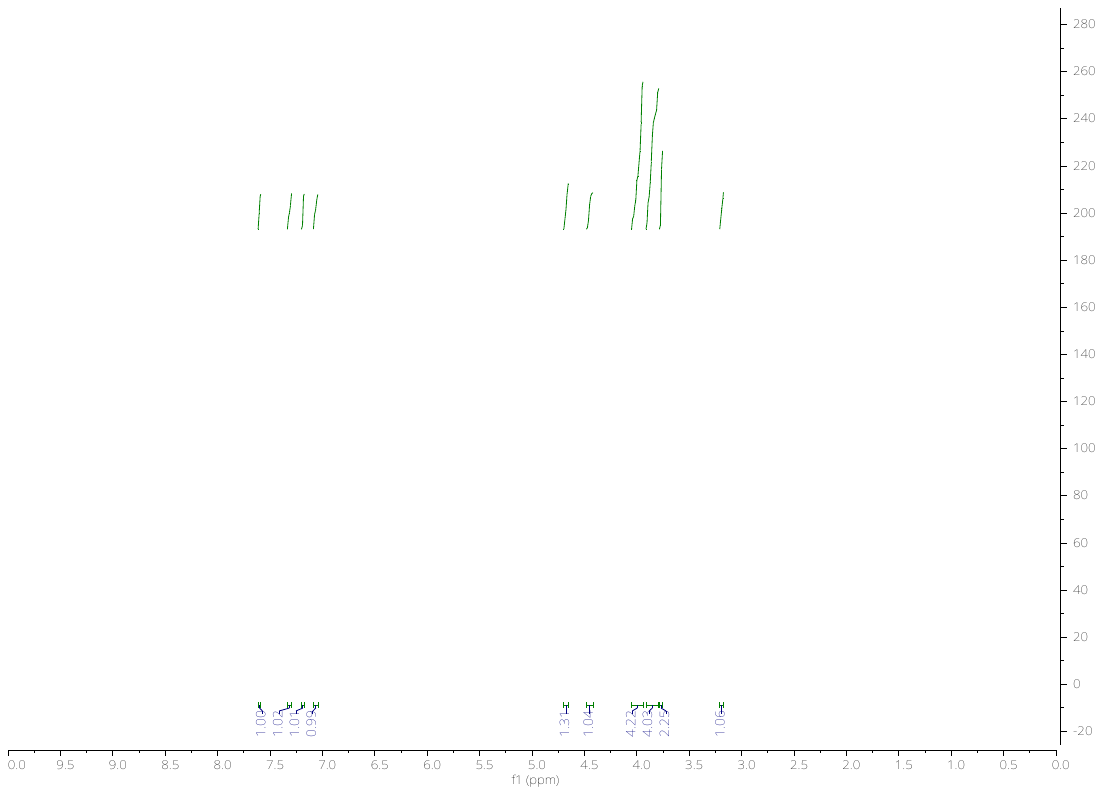

**Figure S3.** ^1^H Spectra of chlorinated G_3_SGW.

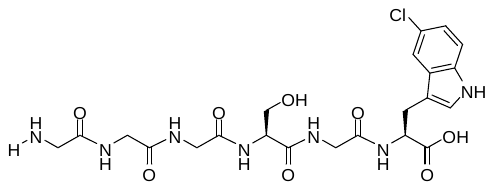

**Figure S4.** Chemical structure of chlorinated G_3_SGW.

1H NMR (400 MHz, CD_3_OD): δ 7.60 (dd, J = 2.1, 0.6 Hz, 1H), 7.31 (dd, J = 8.6, 0.6 Hz, 1H), 7.19 (s, 1H), 7.07 (dd, J = 8.6, 2.0 Hz, 1H), 4.67 (dd, J = 7.8, 5.0 Hz, 1H), 4.46 (t, J = 5.3 Hz, 1H), 4.05 – 3.93 (m, 4H), 3.91 – 3.79 (m, 4H), 3.77 (t, J = 1.2 Hz, 2H), 3.21 – 3.13 (m, 1H).

HRMS (ESI+): m/z calcd for C_22_H_28_ClN_7_O_8_ [M+H]^+^ 554.1761, found 554.1767.

**Table S1.** Amino acid sequence of proteins tested for halogenation showing the N-terminus His-tag in orange, the protein in green, PreScission protease cleavage sequence in blue, FLAG (solubility) tag in red and the HaloTrypTag in black.

| Protein | Sequence | Molecular weight (kD) |
| --- | --- | --- |
| GFP-GGW | MHHHHHHMVSKGEELFTGVVPILVELDGDVNGHKFSVSGEGEGDATYGKLTLKFICTTGKLPVPWPTLVTTLTYGVQCFSRYPDHMKQHDFFKSAMPEGYVQERTIFFKDDGNYKTRAEVKFEGDTLVNRIELKGIDFKEDGNILGHKLEYNYNSHNVYIMADKQKNGIKVNFKIRHNIEDGSVQLADHYQQNTPIGDGPVLLPDNHYLSTQSALSKDPNEKRDHMVLLEFVTAAGITLGMDELYKLEVLFQGPDYKDDDDKGGW | 30.07 |
| GFP-SGW | MHHHHHHMVSKGEELFTGVVPILVELDGDVNGHKFSVSGEGEGDATYGKLTLKFICTTGKLPVPWPTLVTTLTYGVQCFSRYPDHMKQHDFFKSAMPEGYVQERTIFFKDDGNYKTRAEVKFEGDTLVNRIELKGIDFKEDGNILGHKLEYNYNSHNVYIMADKQKNGIKVNFKIRHNIEDGSVQLADHYQQNTPIGDGPVLLPDNHYLSTQSALSKDPNEKRDHMVLLEFVTAAGITLGMDELYKLEVLFQGPDYKDDDDKSGW | 30.10 |
| Stoffel-GGW | MHHHHHHEAPWPPPEGAFVGFVLSRKEPMWADLLALAAARGGRVHRAPEPYKALRDLKEARGLLAKDLSVLALREGLGLPPGDDPMLLAYLLDPSNTTPEGVARRYGGEWTEEAGERAALSERLFANLWGRLEGEERLLWLYREVERPLSAVLAHMEATGVRLDVAYLRALSLEVAEEIARLEAEVFRLAGHPFNLNSRDQLERVLFDELGLPAIGKTEKTGKRSTSAAVLEALREAHPIVEKILQYRELTKLKSTYIDPLPDLIHPRTGRLHTRFNQTATATGRLSSSDPNLQNIPVRTPLGQRIRRAFIAEEGWLLVALDYSQIELRVLAHLSGDENLIRVFQEGRDIHTETASWMFGVPREAVDPLMRRAAKTINFGVLYGMSAHRLSQELAIPYEEAQAFIERYFQSFPKVRAWIEKTLEEGRRRGYVETLFGRRRYVPDLEARVKSVREAAERMAFNMPVQGTAADLMKLAMVKLFPRLEEMGARMLLQVHDELVLEAPKERAEAVARLAKEVMEGVYPLAVPLEVEVGIGEDWLSAKELEVLFQGPDYKDDDDKGGW | 63.67 |
| Stoffel-SGW | MHHHHHHEAPWPPPEGAFVGFVLSRKEPMWADLLALAAARGGRVHRAPEPYKALRDLKEARGLLAKDLSVLALREGLGLPPGDDPMLLAYLLDPSNTTPEGVARRYGGEWTEEAGERAALSERLFANLWGRLEGEERLLWLYREVERPLSAVLAHMEATGVRLDVAYLRALSLEVAEEIARLEAEVFRLAGHPFNLNSRDQLERVLFDELGLPAIGKTEKTGKRSTSAAVLEALREAHPIVEKILQYRELTKLKSTYIDPLPDLIHPRTGRLHTRFNQTATATGRLSSSDPNLQNIPVRTPLGQRIRRAFIAEEGWLLVALDYSQIELRVLAHLSGDENLIRVFQEGRDIHTETASWMFGVPREAVDPLMRRAAKTINFGVLYGMSAHRLSQELAIPYEEAQAFIERYFQSFPKVRAWIEKTLEEGRRRGYVETLFGRRRYVPDLEARVKSVREAAERMAFNMPVQGTAADLMKLAMVKLFPRLEEMGARMLLQVHDELVLEAPKERAEAVARLAKEVMEGVYPLAVPLEVEVGIGEDWLSAKELEVLFQGPDYKDDDDKSGW | 63.70 |
| Spycatcher-GGW | MHHHHHHEMDSATHIKFSKRDEDGKELAGATMELRDSSGKTISTWISDGQVKDFYLYPGKYTFVETAAPDGYEVATAITFTVNEQGQVTVNGKATKGDAHLEVLFQGPDYKDDDDKGGW | 13.26 |
| Spycatcher-SGW | MHHHHHHEMDSATHIKFSKRDEDGKELAGATMELRDSSGKTISTWISDGQVKDFYLYPGKYTFVETAAPDGYEVATAITFTVNEQGQVTVNGKATKGDAHLEVLFQGPDYKDDDDKSGW | 13.29 |
